# Quantitatively Modeling Factors that Influence the Therapeutic Doses of Antibodies

**DOI:** 10.1101/2020.05.08.084095

**Authors:** Yu Tang, Xiaobing Li, Yanguang Cao

**Author notes:** **To whom correspondence should be addressed:** Yanguang Cao, UNC Eshelman School of Pharmacy, UNC at Chapel Hill.

## Abstract

Dose selection and confirmation are critical tasks in the development of therapeutic antibodies. These tasks could become particularly challenging in the absence of robust pharmacodynamics biomarkers or at very flat dose-response curves. Although much knowledge has been acquired in the past decade, it remains uncertain which factors are relevant and how to select doses more rationally. In this study, we developed a quantitative metric, Therapeutic Exposure Affinity Ratio (TEAR), to retrospectively evaluate up to 60 approved antibodies and their therapeutic doses (TDs), and systematically assessed the factors that are relevant to antibody TDs and dose selection patterns. This metric supported us to analyze many factors that are beyond antibody pharmacokinetics and target binding affinity. Our results challenged the traditional perceptions about the importance of target turnovers and target anatomical locations in the selection of TDs, highlighted the relevance of an overlooked factor, antibody mechanisms of action. Overall, this study provided insights into antibody dose selection and confirmation in the development of therapeutic antibodies.

## Introduction

Antibodies have become exceptionally popular and versatile therapeutic agents due to their high target selectivity and long serum half-lives. Therapeutic antibodies have been broadly utilized for the treatment of a variety of diseases, including malignancies and autoimmune, cardiovascular, and infectious diseases.^1, 2^ To date, the US Food and Drug Administration (FDA) has approved more than 80 therapeutic antibodies, and the rate of approval is steadily increasing.^1, 2^

The selection of doses, namely first-in-human doses (FIHDs) in FIH trials, Phase II doses, and therapeutic doses (TDs) in the registration trials, is still one of the most challenging tasks in the development of therapeutic antibodies. The suboptimal selection of doses at each development phase is associated with earlier TD reduction or high drop-out rates in clinical trials, as well as the failure to receive approval.^3^ For conventional small-molecule agents, toxicity-guided dose selection has been routinely applied in FIH trials due to their relatively clear dose-toxicity profiles.^4^ However, dose-toxicity relationships are usually unclear or barely observable for therapeutic antibodies, partially due to their high target selectivity with limited off-target toxicity.^5^ A recent study reported that the dose-limiting toxicity was not observed in more than half of the 82 analyzed FIH trials.^4^ In the absence of clear dose-toxicity profiles, toxicity-guided dose selection strategies may lead to overdosing, increasing the potential risks in late clinical trials.^6^

Given the fact that the conventional toxicity‒guided FIHD selection is not appropriate for many therapeutic antibodies, efficacy has become the primary basis for dose selection.^7^ The minimal anticipated biological effect level (MABEL) has been incorporated in the regulatory agency guideline for choosing FIHDs.^8^ The MABLE approach was first described to select the FIHD by considering the pharmacological activities at the lower end of the dose-response (E-R) curve, such as receptor occupancy (RO) data.^5, 7^ This approach manifests the concept of selecting antibody doses against the expected efficacy rather than the toxicity. Many therapeutic antibodies demonstrated safe and reasonable FIHD selection based on the MABEL approaches.^9^ However, there are no clear guidelines or rationales for antibody dose selection and confirmation in subsequent development phases, resulting in many empirically-based TD selections.^6^ While extensive experience has been gained, the determinants of TDs are still unclear. Little is known about how to determine the TD for an antibody effectively. E-R analyses are normally performed to confirm TDs. However, sometimes the decisions are empirical when the E-R curves are flat, such as several recently approved antibodies.^10-13^

When the long-term efficacy or robust biomarker remains ambiguous in the early stage of clinical trials, RO is frequently adopted as a biomarker to indicate dose adequacy.^5, 14^ RO is typically quantified in circulating cells from peripheral blood samples using flow cytometry, although the peripheral RO (pRO) may not reflect target engagement at the sites of action.^15, 16^ It is still not known how high the pRO must be for a dose to be considered to be therapeutically effective or whether an adequate pRO is related to the mechanism of action. The theoretical analysis based on a target-mediated drug distribution (TMDD) model strengthened the target properties for the selection of TDs. In addition to binding affinity (K_D_), other factors such as target baselines and turnovers, are also believed to be highly involved.^17-20^ Despite these theoretical concepts, the factors that eventually determine the TDs remain elusive and little is known about how to make dose selection efficiently for an antibody under development.

In this study, we applied a retrospective approach to perform a quantitative analysis of up to 60 antibodies that have been approved by the FDA or the European Medicines Agency from 1995 to 2019. To allow comparison across antibody classes, we developed a dimensionless metric, the Therapeutic Exposure Affinity Ratio (TEAR) to (1) systematically evaluate the factors that are associated with the doses and dose selections across antibodies, indications, target properties, and stages of development and (2) identify the potentially overlooked factors. Our analysis provides insights into a critical issue for antibody dose selection and confirmation in the development of therapeutic antibodies.

## Methods

TDs are commonly selected based on E-R relationships. By modeling E-R curves and covariate effects, it is possible to compare the effects across patient populations, and then select TDs with the highest probability of efficacy and tolerable side-effects. However, direct comparisons of E-R relationships across antibodies are not feasible because antibodies have distinct pharmacokinetics, target interactions, mechanisms of action, and target and disease characteristics. Therefore, to evaluate the selected TDs and the influencing factors across diverse classes of antibodies, we developed a quantitative metric: TEAR.

### TEAR ‒ log (C_ss_/K_D_)

pRO is often taken as an intermediate biomarker to reflect the effect of a drug and the adequacy of a dose. The Hill equation is applied to derive pRO based on the target dissociation constant K_D_ and the free drug concentration (C_free_) (Eq.1). In terms of antibody-antigen binding, the equilibrium assumption was made mainly considering that antigen binding is much faster than antibody disposition kinetics. Target binding occurs on a smaller scale than the elimination of antibodies and the typical clinical dosing regimens.

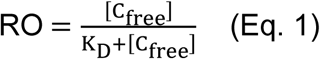

Most approved antibodies are administered at multiple maintenance doses. At maintenance TDs, antibodies achieve steady-state kinetics,

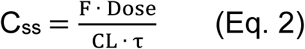

where F is the bioavailability, Dose and τ are the therapeutic maintenance dose and dosing interval, and CL is the antibody systemic clearance at the steady-state concentration (C_ss_). Of note, the constants in Eq. 2 were available for most of the licensed antibodies, allowing for the calculation of C_ss_.

Typically, antibody concentration is much higher than its target concentration; thus, the bound antibody only takes up a small fraction of the total antibody. Therefore, the free antibody concentration approximates the total antibody concentration at steady state, i.e., C_ss_ ≈ C_free_. When target binding occurs in the blood or at anatomical sites in which the antibody can rapidly diffuse into (e.g., lymphoid tissues), the antibody concentration surrounding targets (C_target_) could approximate the concentration in plasma (C_ss_), C_target_ ≈ C_ss_. Integrated into Eq. 1, we derived the pRO as,

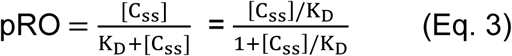

As in Eq. 3, pRO is a function of C_ss_/K_D_. When C_ss_/K_D_ =100 or log (C_ss_/K_D_) = 2, pRO ≈ 99%. If C_ss_/K_D_ > 100 or log (C_ss_/K_D_) > 2, then pRO > 99%. When antibodies with targets in distal tissues, C_target_ is usually less than C_ss_, which entails a higher C_ss_ for an adequate C_target_ and a higher pRO for treatment efficacy.

The log-transformed C_ss_/K_D_ values fall within a normal distribution, which enables robust statistics. TEAR is thus defined as log (C_ss_/K_D_) to reflect the relationship between antibody therapeutic concentration and target binding affinity. Notably, TEAR considers the variance associated with F, CL, and K_D_. The remaining difference in TEAR would reveal the variance related to other factors, such as the target location and turnover rates. Therefore, TEAR supports the examination of the influence of indications, target locations, target baselines and turnovers, mechanisms of action, and all other factors on the TDs of antibodies.

### Categorization of the Target locations and Properties

#### Circulation vs. tissue

We categorized antibodies with targets that are predominantly expressed in either the blood or the lymph system or on circulating lymphocytes as the circulation group, for which plasma concentrations should approximate the antibody. For example, CD20 is a membrane-associated receptor that is widely expressed on B-lymphocytes. We categorized CD20 into the circulation group because of the location of CD20-expressing cells. Antibodies with targets expressed in tissues, such as skin and tumors, were included in the tissue group. We included antibodies with targets that are present in both the circulation and tissues in both groups. Notably, the distribution of targets is considered under pathological conditions.

#### Soluble vs. membranous targets

We categorized the receptors that are expressed on the cell membrane as membranous targets, such as epidermal growth factor receptor (EGFR). The ligands that freely distributed in the blood, the lymphatic system, and interstitial fluids were defined as soluble targets, such as viruses and cytokines. If a target has both soluble and membranous forms, it was included in both categories. The shredding of membranous targets was not considered in this study.

#### Specific anatomical sites

According to the National Cancer Institute criteria, we divided antibodies in treatment of cancers into the following groups: the blood and lymph system, the breast and digestive/gastrointestinal (GI) system, the gynecological system, the head and neck, the musculoskeletal system, the neural system, the respiratory/thoracic system, and the skin. The anatomical sites were also specified for the antibodies in the treatment of autoimmune diseases, such as antibodies acting in the blood and the lymph system (i.e., IL-1β, IL-4R, IFNγ, IgE, CD80/CD86, BAFF, α4β1, CD20, IL-6, and IL-6R), synovial fluid (i.e., CD80/CD86, IL-1β, BAFF, CD20, IL-6R, IL-6, and TNF-α), bronchoalveolar lavage fluid (i.e., IL-5 and IL-5R), cerebrospinal fluid (CD20 and α4β1), or skin lesions (i.e., IL-17R, IL-17, and IL-23).

### Target Turnover Rates

The target turnover rate is calculated by multiplying the target baseline concentration (nM) with the target degradation rate (hr ^-1^):

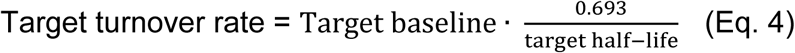

The target baselines and degradation rates or half-lives were collected from literature. For a soluble target, the plasma baseline was directly adopted, and the degradation rate was derived based on reported plasma half-life. For a membranous target, the degradation rate was assumed to be equal to the antibody-complex endocytosis rate, but the target baseline is typically not available. Thus, an equivalent plasma concentration was calculated using the following equation.

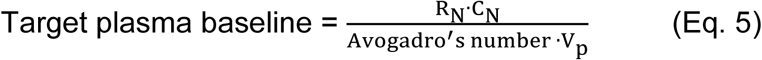

R_N_ is the receptor number/cell. C_N_ is the cell number per 1 *μ*L blood or 1 *μ*g tissue. For tumors, 1 *μ*g is close to 10^9^ cells, the detection limit of most tumor types.^21^ V_p_ represents the plasma volume (50 mL/kg).^22^ Of note, the target turnover rate reflects the total turnover rate in the system, which enables direct comparisons across target types and locations.

### Dose Selections in FIH and Phase II trials

#### FIHDs, maximum administered doses (MADs), and Phase II doses

When available, the first-tested dose in the FIH trial of a given antibody was defined as the FIHDs. The highest tested doses in the FIH trials were defined as the MADs. All Phase II doses recorded by the FDA reviews were included in this study.

#### Metrics for analyzing single-dosed antibodies in FIH trials

For the antibodies that only have single dose FIHDs or MADs in the FIH trials, we applied the log (C_FIHD_/K_D_) and log (C_MAD_/K_D_) to evaluate factors that influence the FIHDs and MADs. These metrics are similar to the TEAR, but more relative to the MABEL approach. The equations for calculating these two MABEL-related metrics were shown below.

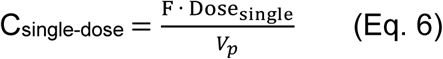

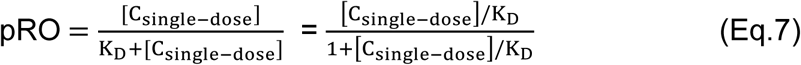

#### Phase II selection rationales

MABEL approach-based biomarkers included the experimentally-acquired or model-informed target engagement for the circulating cell surface targets, the saturation of target-mediated clearance, and the pre-defined downstream pharmacological responses. Efficacy‒based biomarkers include pharmacological responses and clinical outcomes that are not directly associated with target engagement.

## Results

### Most Approved Antibodies Have TDs Oversaturating the Peripheral Targets

As shown in Eq. 3, when TEAR = 2, pRO ≈ 99%, indicating that the circulating targets were almost completely saturated at the TDs. As seen in Figure 1, the TEARs of 60 approved antibodies ranged from 1.1 to 5.9, which yielded approximately 92% to 100% pRO. Most of the surveyed antibodies (55 out of 60) had TEARs > 2, suggesting that most antibodies had almost completely saturated pROs at their TDs. Moreover, 39 of those 55 antibodies had TDs that are at least 10-time greater than the doses yielding 99% pRO, and 18 antibodies had TDs that are at least 100‒times higher than doses yielding 99% pRO. Unexpectedly, some antibodies with extremely high TEARs > 4, such as ravalizumab and emapalumab, have their targets primarily in the circulation, suggesting that the TDs for those antibodies are not just influenced by their CL and K_D_, or solely explained by their target location. More evaluations are warranted.

**Figure 1.**
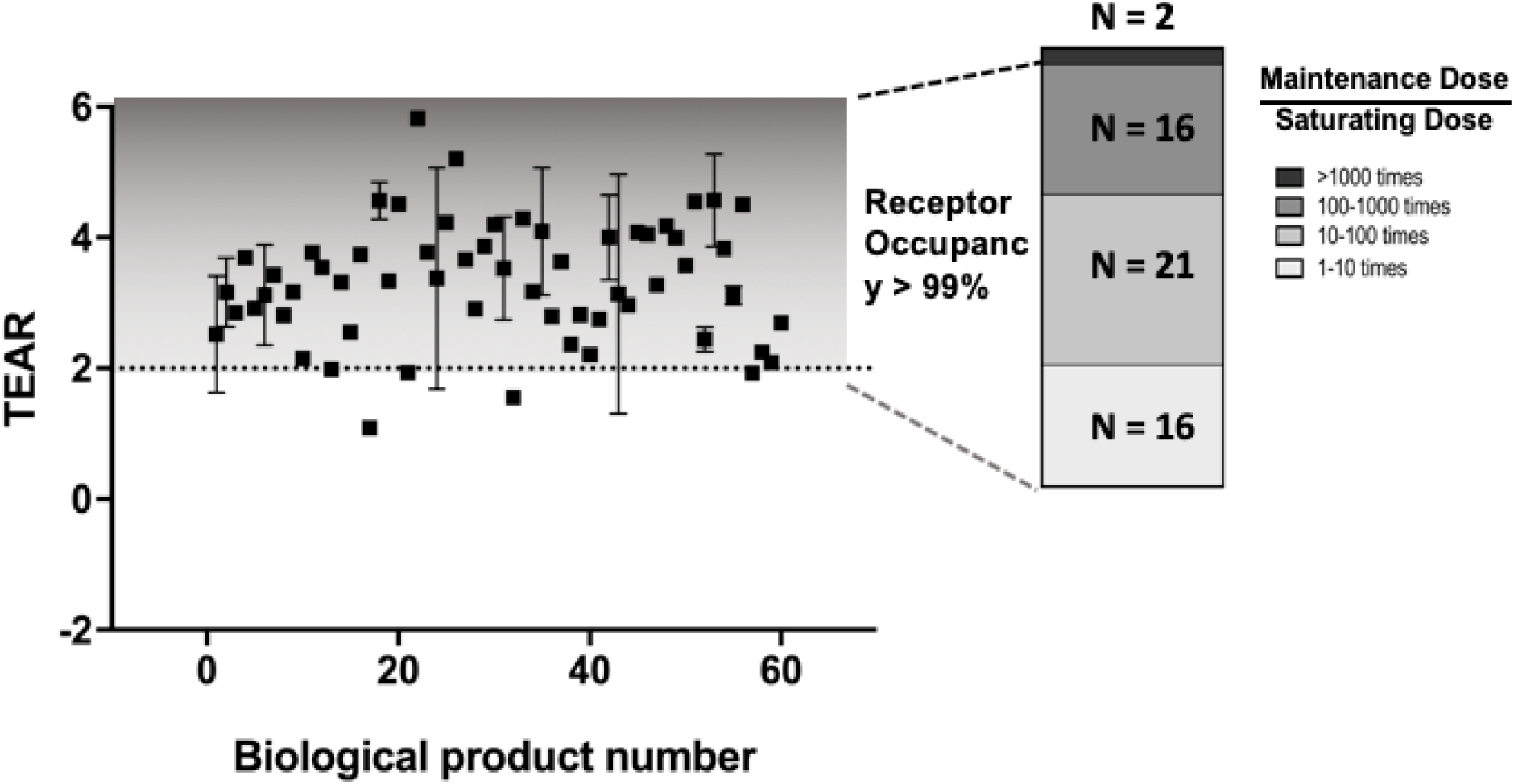
Most approved antibodies have therapeutic doses (TDs) oversaturating targets in plasma, shown by the TEAR metric. Up to 60 licensed antibodies were included in the analysis. Most of the surveyed antibodies had TEARs > 2, indicating those antibodies could saturate their targets in peripheral blood (peripheral receptor occupancy [pRO] ≈ 99%) at the TDs. As shown in the bar chart, two antibodies had TDs >1000-fold, 16 had TDs >100-fold, 21 had TDs 10 ~ 100-fold, and 16 had TDs 1 to 10-times higher than the doses yielding nearly saturated pRO. Each dot represents the TEAR of an antibody in mean ± SD. The antibodies are numbered in alphabetical order and shown on the x-axis.

### Effect of Target Locations, Forms, and Turnovers on TDs

Theoretical modeling suggested that target baselines and turnovers both significantly influenced the TDs of antibodies. We calculated the turnover rates of all the targets with at least one approved antibody. The target turnover rate is the target baseline multiplied by the degradation rate (Eq. 4) reflecting the total replenishing target per unit time. A total of 52 antibodies, along with turnover rates of their targets, were summarized in **Supplementary Table 2**. The correlation between target turnover rates and TEARs was depicted in Figure 2. No significant correlation was detected between the TEARs and target turnovers for the surveyed antibodies (P = 0.26, Pearson’s correlation). This observation contradicted the previous theoretical analysis, suggesting that the influence of target turnover on the selection of TDs is probably not as straightforward as we generally thought, and more investigations are warranted

**Figure 2.**
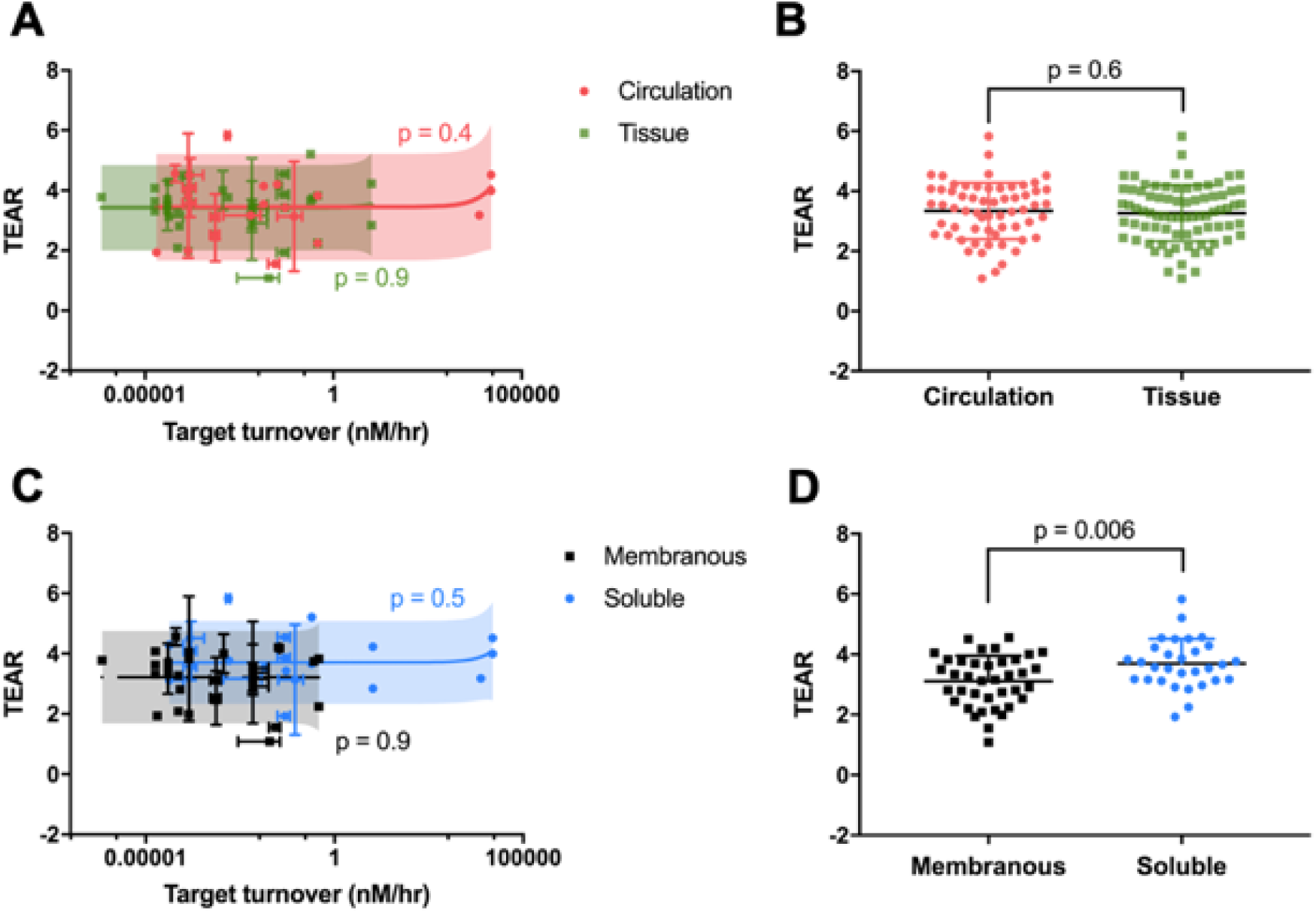
Effects of target anatomical locations, forms, and turnovers on therapeutic doses (TDs). **(A)** Target turnovers are not relevant to antibody TDs, either in the circulation group or the tissue group (P = 0.4, P = 0.9, respectively, Pearson’s correlation). Dots represent the mean values. Horizontal bars represent SD in the target turnover rates. Vertical bars represent SD in the TEARs. The red and green shadows represent the 90% prediction intervals. **(B)** Target anatomical location does not have a significant impact on TDs. There is no significant difference between the TEARS in the circulation and tissue groups (P = 0.6, unpaired Student’s *t*-test). Each dot represents the mean TEAR of an antibody. The bars represent mean ± SD values. **(C)** Target turnover is not a significant factor to TDs in both the membranous and soluble groups. TEARs have no significant correlation with target turnovers, regardless of the solubility of the target (P = 0.9, membranous targets; P = 0.5, soluble targets. Pearson’s correlation). Each dot represents the mean value of an antibody. **(D)** The TEARs are significantly different between the soluble and membranous groups (P = 0.006, unpaired Student’s *t*-test).

To further elucidate this, we divided the antibodies into two groups based on their target anatomical locations: circulation vs. tissue. Still, no statistical correlation was observed between the TEARs and target turnovers in either circulation or tissue group (P = 0.4 vs. P = 0.9, Pearson’s correlation**, Figure 2A**). For antibodies with targets outside the circulatory system, we typically expect a relatively high dose to provide adequate target exposure. However, we did not find a statistical difference between the two groups of antibodies with two distinct target locations (P = 0.6, unpaired Student’s *t*-test, Figure 2B).

The antibodies were further divided into two groups based on their target solubility: membranous vs. soluble targets. There was still no significant correlation detected between the TEARs and target turnovers in either membranous or soluble group (P = 0.5 vs. P = 0.7, Pearson’s correlation, Figure 2C). However, we found TEARs is notably higher for the antibodies with soluble targets compared to those with membranous targets (3.7 vs. 3.1, P = 0.006, unpaired Student’s *t*-test, Figure 2D). This observation suggested that a higher antibody TD is usually needed to effectively suppress the soluble target.

Antibodies with extremely high or low target turnovers were specifically examined. Some antibodies had both high target turnovers and high TEARs, such as eculizumab. The target of eculizumab (i.e., complement component 5 [C5]) had the highest target turnovers (15,384 nM∙hr^-1^) among all the analyzed targets. Eculizumab was approved with a TEAR as high as 4.52, which was higher than 90% of the analyzed antibodies (**Supplementary Table 1**). Compared to C5, the target of risankizumab (i.e., interleukin 23 [IL-23]) had a significantly lower target turnover (0.0482 nM∙hr^-1^, **Supplementary Table 1**). However, risankizumab was approved with an even higher dose, resulting a higher TEAR than eculizumab (4.55 vs. 4.52, **Supplementary Table. 1**). Another striking case is erenumab, an antibody targeting calcitonin gene-related peptide receptor (CGRPR). CGRPR has the slowest turnover (7×10^-7^ nM∙hr-1) among the investigated targets. However, erenumab has a TEAR of 3.78, which is higher than 70% of the surveyed antibodies. Overall, while the target turnover probably influences the TDs of some antibodies, its general relevancy to the selection of TDs of antibodies warrants further investigation.

### Disease Types and Sites of Action Are Not Significantly Relevant to TDs

To further explore the factors, we cross-examined the TEARs in four disease-target scenarios: circulation-soluble, circulation-membranous, tissue-soluble, and tissue-membranous. As shown in Figure 3A, the common diseases in each scenario were autoimmune diseases with targets in the circulatory system (CS), hematologic malignancies (CM), autoimmune diseases with tissue targets (TS), and solid tumors (TM). Of note, the CS group included antibodies with targets such as BAFF, IL-1β, IFNγ, IL-5, IgE, IL-6, and IL-6R. The CM group included antibodies with targets primarily expressed in the circulating cells, such as PD-1, CD38, SLAMF7, and CD20, which are mostly related to immune cell functions. As seen in Figure 3B, the ranking of TEARs for each group were: CS > TS > TM > CM (mean TEAR = 3.8, 3.5, 3.0, and 2.5, respectively). The TEARs were significantly lower for the antibodies in the CM group than in the CS group (P = 0.001, unpaired Student’s *t*-test, Figure 3B). No significant difference was detected between any other two groups. Moreover, no significant correlation between the TEARs and the target turnovers was observed in any of the four groups (**Supplementary Figure 1**).

**Figure 3.**
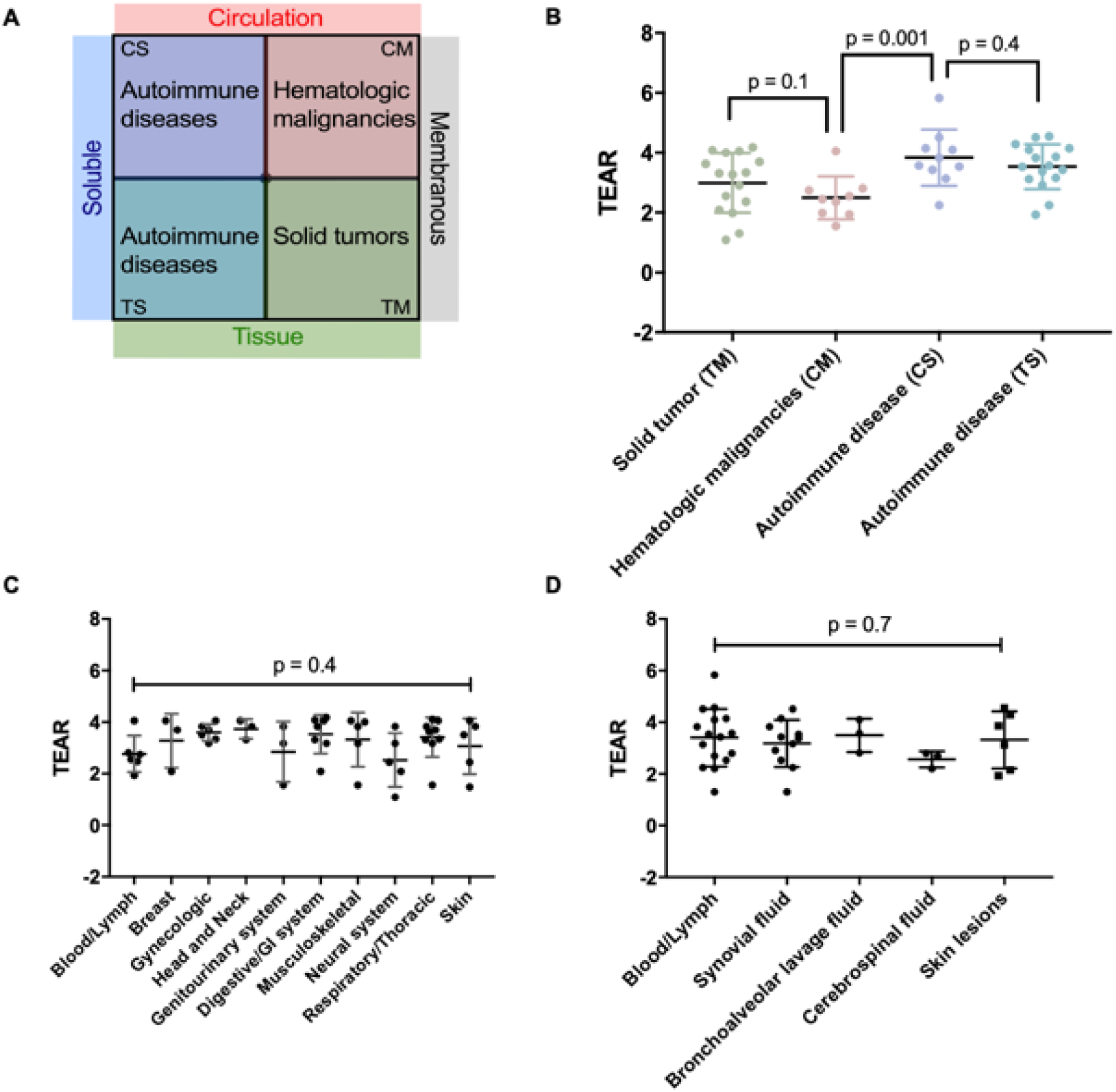
The relevancy of indications, and target tissues to to therapeutic doses (TDs). **(A)** The antibodies were categorized into four disease-target scenarios. The major diseases in each scenario are autoimmune diseases with targets in circulation (circulation-soluble, [CS]), hematologic malignancies (circulation-membranous, [CM]); autoimmune diseases with tissue targets (tissue-soluble, [TS]); and solid tumors (tissue-membranous, [TM]). **(B)** A significant difference was only observed between the hematologic malignancies group and the autoimmune diseases with tissue target group (P = 0.001, unpaired Student’s *t*-test). No difference in the TEARs was observed in the other groups. **(C)** The TEARs are not significantly different between the tumor anatomical locations (P = 0.4, ordinary one-way ANOVA). **(D)** The TEARs are not significantly different between the target anatomical locations in autoimmune diseases (P = 0.7, ordinary one-way ANOVA). Each dot represents the TEAR of an antibody. The data is represented in mean ± SD.

The target anatomical locations were further explored. The TEARs across tumor types were compared. No significant influence of tumor types on TEARs was observed (P = 0.4, one-way ANOVA, Figure 3C). A similar analysis for antibodies in treatment of autoimmune diseases suggested that the TEARs were not substantially different across anatomical sites (P = 0.7, ordinary one-way ANOVA, Figure 3D). Collectively, these observations suggested that disease types and target anatomical locations have moderate relevance to the selections of TDs.

### Mechanism of Action is a Pivotal Factor in TDs

In general, therapeutic antibodies elicit efficacy via three distinct mechanisms (Figure 4A): neutralizing soluble targets (soluble target neutralization, [STN]), binding to membranous targets to suppress intracellular singling (membranous signaling suppression, [MSS]), and triggering immune functions to lyse targeted cells (immunomodulatory function, [IF]).Antibodies in the STN group were mostly from the CS and TS groups (Figure 3), while the CM group primarily consisted of antibodies with mechanisms of MSS and IF (Figure 4A). The major action of the antibodies in the STN group is to neutralize the pathological factors and then alleviate inflammation or malignancies. The antibodies in the MSS group bind to a membranous receptor to suppress or trigger intracellular signaling pathways to inhibit cell proliferation and stimulate cell death. In the IF group, antibodies can elicit efficacy by triggering effector cell immune functions or recovering the inhibition of immune cell functions. For the antibodies that eradicate target cells via effector functions, their Fc domains engage with the Fcγ receptors on the effector cells upon antibody: antigen binding, activating the effector cells to lyse the target-expressing malignant cells Of note, antibodies in the CM and CS groups may have distinct mechanisms of action. In the CM group, most antibodies elicit their treatment effect via mechanisms of MSS and IFs, while the antibodies in the CS group primarily work through the mechanism of STN.

**Figure 4.**
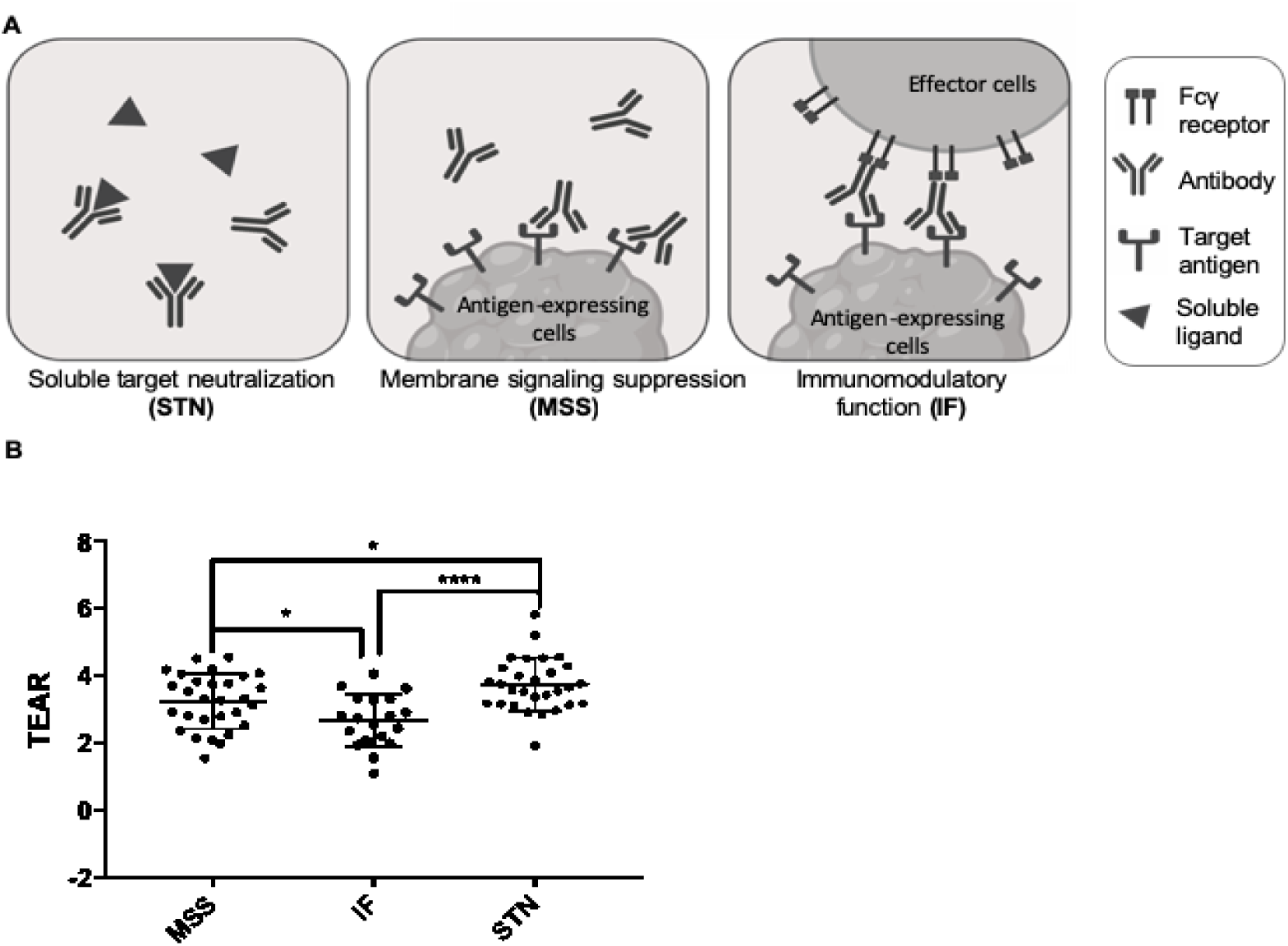
The mechanism of action is a pivotal factor in discerning therapeutic doses (TDs). **(A)** Antibodies elicit therapeutic efficacy via three distinct mechanisms: soluble target neutralizing (STN), membranous signaling suppression (MSS), and immunomodulatory function 14. **(B)** The TEARs are significantly different between antibodies with varying mechanisms of action. The TEARs in the MSS and STN groups are significantly higher than the IF group (P = 0.02, P < 0.0001, unpaired Student’s *t*-test). Each dot represents the TEAR of an antibody. The data is represented in mean ± SD.

We compared the TEARs across three groups of antibodies based on their mechanisms of action. Interestingly, there were significant differences in the TEARs between each two groups (Figure 4B). The antibodies in the IF group had the lowest TEARs (mean TEAR = 2.6), whereas the antibodies that act through STN appeared to have the highest TDs (mean TEAR = 3.7). The antibodies in the MSS group had significantly lower TEARs in comparison to the ones in the STN group (3.2 vs. 3.7, P = 0.02, unpaired Student’s *t*-test, Figure 4B). The antibodies in both MSS and STN groups had significantly higher TEARs than the antibodies in the IF group (P = 0.02, P < 0.0001, unpaired Student’s *t*-test). Among the five antibodies with TEARs < 2, four antibodies act by modulating immune functions, i.e., from the group of IF. For instance, the anti-GD2 antibody dinutuximab, which binds to neuroblastoma cell surface GD2 and induces cell lysis via effector functions, had the lowest TEAR among the 60 surveyed antibodies (TEAR = 1.08, **Supplementary Table 1**). While panitumumab mainly eradicates EGFR-expressing cells via the mechanism of MSS, necitumumab and cetuximab elicit tumor-cell killing effects by both mechanisms of MSS and IFs. Cetuximab and necitumumab had lower TEARs than panitumumab (3.3 vs. 4.1, 3.6 vs. 4.1, respectively. **Supplementary Table 1).** These findings strongly indicated the influence of the mechanism of action on the TDs of antibodies.

### Mechanism of Action Also Has Impacts on FIHD and Dose Escalation

To evaluate doses in FIH and Phase II trials and the dose selection rationales, we applied two metrics that were similar to TEAR, i.e., log (C_FIHD_/K_D_) and log (C_MAD_/K_D_), to support comparison across antibodies with distinct pharmacokinetic properties and target affinities. Among the 40 antibodies that were analyzed, we found high variabilities in the FIHDs and the dose escalation ranges (DERs) in their FIH trials. As shown in Figure 5, the lowest log (C_FIHD_/K_D_) was 10,000-fold lower than the highest one. With regard to the DERs, while the MADs of some antibodies were only 2-times higher than the FIHDs, other antibodies had up to 10,000 DERs. Significant variations were also noticed in the Phase II dose selections. Some antibodies, such as atezolizumab and avelumab, only had a single dose tested in Phase II trials; many other antibodies had multiple doses tested or went another round of dose escalation in the Phase II trials. For most antibodies, the Phase II doses were selected from the DERs in the FIH trials, and almost all of the surveyed TDs were from the Phase II-tested doses.

**Figure 5.**
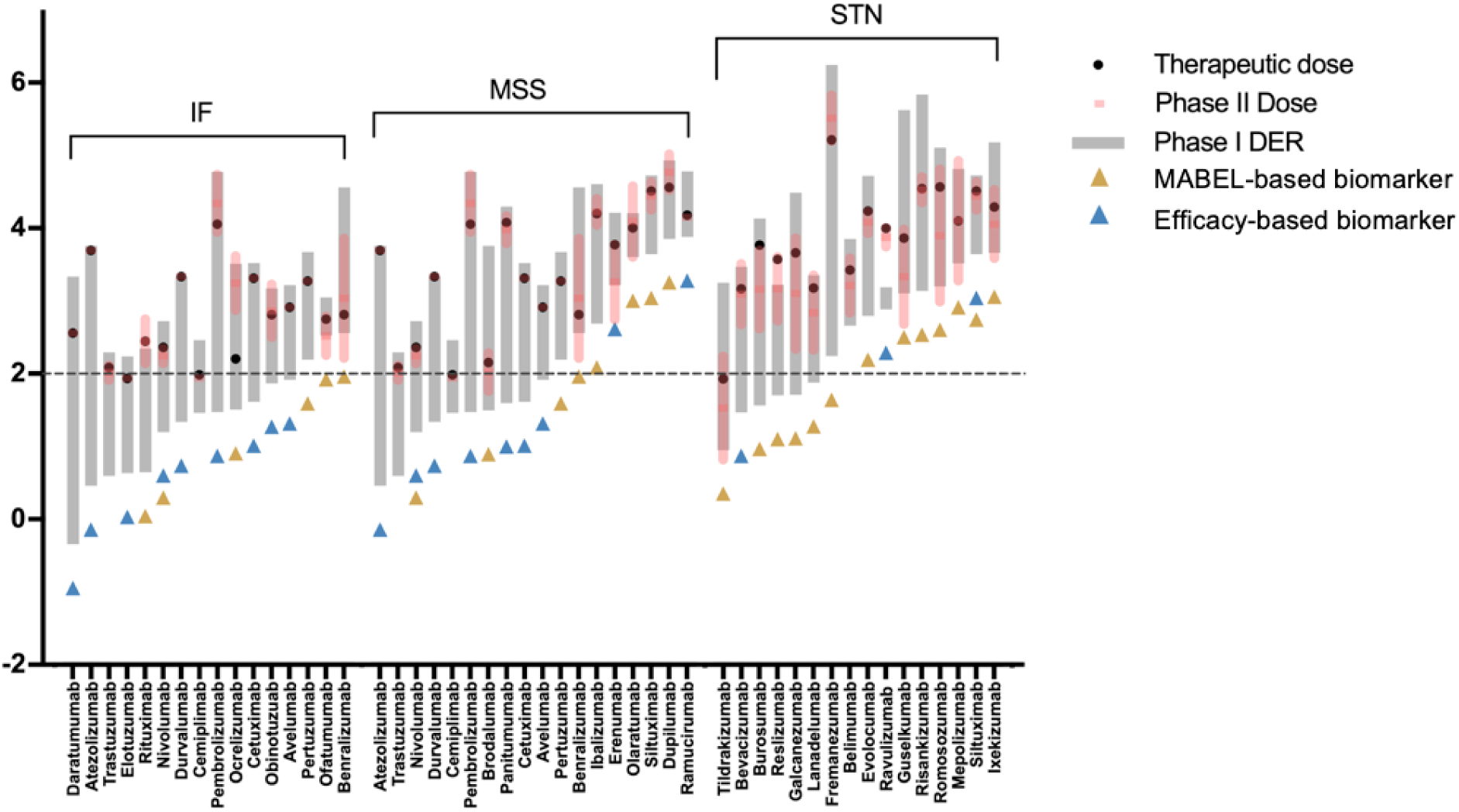
Mechanism of action has an impact on antibody dose escalations. The dose-escalating ranges (DERs) in first-in-human (FIH) trials are denoted by the grey bars, the lower and upper edges of which represent the FIH doses and maximum-administered doses (MADs) in FIH trials. The TEARs of the Phase II doses are given in mean ± SD, demonstrated by red bars and shadows. The solid black circles represent the TEARs of therapeutic doses (TDs). The Blue and yellow triangle symbols denote the antibodies that used MABEL approaches or treatment efficacy as the Phase II dose selection rationale. Forty antibodies were included in the analysis and grouped by their mechanisms of action. The names of the tested antibodies are indicated on the x-axis.

Notably, the mechanism of action also has an impact on the FIHDs and the subsequent Phase II dose selections. More than 80% of the therapeutic antibodies that act through effector functions (in the IF group) had relatively low and conservative FIHDs, and their log (C_FIHD_/K_D_) values were all < 2, significantly lower than the ones in the MSS and STN groups (1.4 vs. 2.2, P = 0.02; 1.4 vs. 2.5, P = 0.005. unpaired Student’s *t*-test. **Supplementary Figure 3A**). We also evaluated other factors that may influence FIHD and dose escalation. The log (C_FIHD_/K_D_) values were slightly lower in first-in-class antibodies in comparison to next-in-class antibodies. However, no statistically significant difference was detected between the two groups (2.0 vs. 2.2, P = 0.5, **Supplementary Figure 3B**). Target properties, including target locations, baselines, and degradation rates, were not found to have a significant impact on dose selections (**Supplementary Figure 3C ‒ F**).

We found that 92% of the Phase II TEARs were > 2, indicating that almost all the Phase II doses were adequate to saturate the peripheral targets (Figure 5). We also found that Phase II dose selection differed across antibodies with distinct mechanisms of action. About half of the antibodies in the IF group only had one dose tested in the Phase II trials, around 33% of antibodies in the MSS group were tested at a single dose, but none of the antibodies in the STN group had single dose in Phase II trials. For the antibodies with multiple doses tested in Phase II trials, the ranges of doses were higher in the STN group in comparison to antibodies in the IF and MSS groups (Figure 5). The rationales for Phase II dose selection were also different across the mechanisms of action. Most of the antibodies in the IF group and half of the antibodies in the MSS group adopted MABEL‒ based biomarkers when selecting Phase II doses. Conversely, efficacy-based biomarkers were frequently used to select Phase II doses in the STN group.

## Discussions

Dose selection is critical for a therapeutic antibody, especially at its first clinical entry and the following treatment outcome optimization. However, the current antibody dose selection approaches are still empirical.^6^ The importance of the underlying physiological and pharmacological factors beyond clearance and target affinity in dose selection remains unknown. A retrospective examination of the dose selections of licensed antibodies will help us identify the potential factors that influence dose selection. However, direct investigations and comparisons of antibody doses are greatly biased by high variabilities in antibody administration routes, dosing intervals, clearances, and binding affinities. Because it is not feasible to directly compare treatment efficacy across antibodies, pRO has become an intermediate biomarker for evaluating dose adequacy.^5, 14^ In this study, we used a dimensionless quantification metric, TEAR, to (1) perform direct evaluations of doses across antibodies after normalizing the differences in antibody bioavailability, dosing interval, clearance, and affinity, (2) explore the pRO of the licensed antibodies at their tested doses and TDs, from the first clinical entry to dose confirmation, and 3) evaluate the critical factors that influence dose selections at each development stage, especially to the confirmation of TDs. TEAR is a metric that directly reflects pRO. A TEAR threshold of 2 represents a nearly complete pRO (pRO ≈ 99%). A TEAR > 2 indicates that the evaluated dose is more than adequate to saturate the peripheral targets, implying that additional factors are involved in high TDs.

In our analysis, one striking observation was that the target baseline and turnover were not apparently related to the antibody TDs (Figure 2, **Supplementary >Figure 1 and 2**), which contradicted previous theoretical predictions. The contradiction could be due to multiple confounding factors in the selections of TDs. Previous theoretical work evaluated the factors largely based on the standard TMDD model, where the target anatomical locations, antibody mechanisms of action, and many other pharmacologically relevant factors were not considered.^18, 20^ Furthermore, those theoretical simulations were often performed by shifting factors one-by-one, without appreciations of the interactions and correlations. For instance, antibodies with targets in solid tumors usually elicit actions by triggering effector functions, which may require a relatively lower TD and partial receptor engagement for therapeutic efficacy.

Our study showed that the primary anatomical location of targets was not a significant predictor to antibody TDs. Antibodies, due to large molecular sizes, typically have poor tissue penetration;^23^ thus, antibodies with targets beyond the vascular space or the lymphoid system are expected to have high TDs. However, our analyses did not find strong evidence to support the influence of target location on the selection of TDs. In fact, the therapeutic values of local vs. systemic antibody actions are still unclear for several classes of antibodies. Many antibodies have their targets present throughout the system. The local tissue distribution of these antibodies may be not directly related to the overall therapeutic effect or the selection of TDs. For example, for antibodies in treatment of metastatic malignancies, the therapeutic efficacy often goes beyond regional drug action and the regional drug distribution. One recent study showed that metastases at multiple anatomical sites responded similarly to pembrolizumab, even though these tissues have varying antibody permeabilities and distributions.^24^ This observation was echoed by other studies indicating the critical therapeutic values of systemic immune response.^25-27^ The efficacy, beyond the primary target locations, at the surrounding tissues and other lymphoid tissues has also been highlighted to the success of treatment.^28-30^ Similarly, for cytokine-neutralizing antibodies in the treatment of autoimmune diseases, whether or not the local injection of antibodies could yield a better effect than the systemic administration has been a long-standing debate.^31, 32^ The immune system is very dynamic. Constant exchanges and communications occur across anatomical locations. It is likely that the efficacy of many antibodies is largely defined by the actions at the systemic level. Thus, the required TDs are not constrained by the primary target anatomical locations. We recently used a novel imaging approach and observed that ROs inside tumors were never complete even at a supratherapeutic dose of antibody that was 5-times higher than the therapeutic dose.^33^ This further implies that adequate exposure of antibodies to the primary anatomical site may not be feasible or critical to therapeutic efficacy. For such antibodies, the TDs might be largely defined by their actions at the systemic level, which undermines the impact of antibody distribution into the primary target site to the selection of TDs.

The mechanisms of action for the selection of TDs were crucial based on our analysis. Among all the antibodies applied in cancer treatment (**Supplementary Table 1**), those in the IF group have the lowest TEARs in comparison to the antibodies with the other mechanisms of action (Figure 4B). This finding highlighted that relatively lower doses or partial target engagement can sufficiently activate effector, which make the host immune functions become critical for antibodies in the IF group.^34^ For example, in patients with squamous cell carcinoma treated with cetuximab, survival was much higher in individuals with stronger immune responses ^35^. Thus, quantitative analyses of the immune functions during antibody treatment are beneficial to establishing dose-response relationships. Many studies have highlighted the values of understanding the dynamic interactions between the tumor and the immune system to understand antibody efficacy and resistance.^36^ Despite the importance of immune functions, yet, a quantitative relationship between dose, effector functions, and treatment efficacy has not been identified for most antibodies. The unclear relationship is partially because the effector functions are very dynamic and challenging to quantify, obscuring the role of effector functions in the net tumor-killing effect. Sensitive biomarkers for effector functions and quantitative modeling approaches are highly desirable for understanding effector functions in antibody-based treatment and valuable for establishing dose-response relationships.

We also observed that antibody FIHDs and dose-escalating patterns differed greatly between the antibodies with distinct mechanisms of action. Most antibodies in the IF group had affinity-normalized concentration, log (C_FIHD_/K_D_) < 2. This finding corresponds to a recent FDA report, which showed that more than half of the immunomodulatory antibodies in investigational new drugs (INDs) were far from saturating their targets at FIHDs.^4^ The conservative FIHD selection was due to less-certain dose-response relationships for the antibodies that could modulate immune functions. After the lessons learned from the TGN1412 case,^37^ the MABEL approach is recommended to prevent potential adverse events in which low FIHDs that result in minimum pRO (~ 10%) are typically chosen in FIH trials.^5^ Surprisingly, the normalized FIHDs between first-in-class and next-in-class antibodies were not significantly different. The Phase II dose selection patterns were also associated with the antibody’s mechanisms of action. The antibodies in the IF and MSS groups were more likely to have fewer Phase II doses tested than the antibodies in the STN group, which probably reflects the different indications of those antibodies. As known, the antibodies with IF and MSS functions are often applied in oncology, where fewer but effective doses are preferable when examining the treatment efficacy.^38^ Most antibodies in the STN group are for autoimmune disease treatment in which identifying the optimal dose-response relationships are more desirable in Phase II trials. The different Phase II dose selection goals may also explain why most of the antibodies in the STN group used efficacy-based biomarkers to select the Phase II doses, in contrast to the commonly applied MABEL approach for antibodies in the MSS and IF groups.

Our analyses have a few limitations. First, we applied target affinity (K_D_) values measured under *in vitro* conditions to derive TEAR. Even though we considered an average value of multiple measurements, sometimes, there was a gap between the *in vitro* and physiological binding conditions, where many stromal factors sterically hinder an antibody’s interactions with its targets. Second, the turnovers for targets in solid tumors were calculated with an assumption that only 10^9^ cells were present.^21^ It is known that the number of tumor cells largely varies across cancer patients at different stages of diagnosis. Third, we only analyzed pharmacologically relevant factors. As known, decisions about antibody doses are often made after considering many practical and pharmacoeconomic factors, such as ethical restrictions, cost-effectiveness relationships under different dosing regimens.

In conclusion, our study offered a quantitative method to support direct and systematic evaluation of antibody doses and dose selection patterns. Our analyses provided some insights into the factors that could have an impact on antibody efficacy and the selections of TDs. Our results highlighted the importance of the mechanism of action and challenged the traditional perceptions about the importance of target turnovers and locations in the selection of TDs. Thus, this study has strong implications for the future development of therapeutic antibodies.

## Study Highlights

### What is the current knowledge on the topic?

Antibodies have been the most popular therapeutic agents for the treatment of a variety of diseases. To date, more than 80 antibodies have been approved by the US Food and Drug Administration (FDA), and the yearly approval number is increasing. The selection of an appropriate human starting dose and the confirmation of a therapeutic dose (TD) have been a long-standing issue in the development of antibodies. However, the current approaches for antibody dose selection remain largely empirical.

### What question did this study address?

How can we systematically evaluate the TDs of the approved antibodies and quantitatively identify the influencing factors in the process of dose selection and confirmation?

### What does this study add to our knowledge?

Our results challenge the traditional perception that antibodies should have high TDs when the target has a high baseline, rapid turnover, and deep anatomical location.

Instead, we highlighted the mechanism of action, which is an overlooked factor, in the selection of FIHD and confirmation of TDs.

### How might this change clinical pharmacology or translational science?

This study provided insights into a critical issue with respect to antibody dose selection and confirmation in the development of therapeutic antibodies.

