## Supplementary Material for "Quantitatively Modeling Factors that Influence the Therapeutic Doses of Antibodies"

### Supplementary Materials

#### Data Sources

In total, 60 antibodies and antibody-based biologics that were approved by the EMA or FDA between 1995 and 2019 were included in our analyses. The excluded antibodies were: (1) the withdrawn or marketing discontinued antibodies or fusion proteins (e.g., muromonab-CD3, efalizumab, tositumomab-I131, daclizumab, catumaxomab, edrecolomab, Nebacumab, and alefacept); (2) antibody-drug conjugates (e.g., brentuximab vedotin, ado-trastuzumab emtansine, inotuzumab ozogamicin, gemtuzumab ozogamicin, and polatuzumab vedotin); (3) bi-specific antibodies (e.g., emicizumab and blinatumomab); (4) antibodies with neither F CL, nor $K_D$ information available (e.g., Ibritumomab tiuxetan, efmoroctocog-α, eftrenonacog-α, mogamulizumab, and ustekinumab); (5) single-dosed antibodies that cannot support $C_{ss}$ calculation, including abciximab, alemtuzumab, basiliximmab, bezlotoxumab, idarucizumab, obiloxaximab, and raxibacumab; and (6) local dosing and acting antibodies (e.g., aflibercept, ranibizumab, and brolucizumab).

We obtained the labeled TDs and dosing regimens from the Drugs@FDA website and the literature. The following information was also collected: (1) molecular weight, (2) administration route, (3) bioavailability (F), (4) *in vitro* antigen-binding dissociation constant ($K_D$), (5) the antibody systemic clearance (CL), (6) target abundance and half-life, (7) the approved indications, and (8) the mechanism of action. More specifications for these parameters were provided in the **Supplementary Table 1**.

**Supplementary Figure 1**

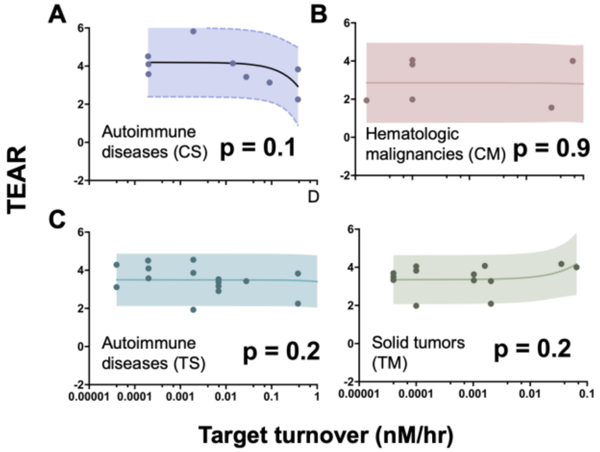


**Supplementary Figure 1. Target turnovers are not relevant to TEARs in four disease-target scenarios.** Dots represent the mean values. Horizontal bars represent SD in the target turnover rates. Vertical bars represent SD in the TEARs. The shadows represent the 90% prediction intervals. No significant correlation between target turnovers and TEARs was observed in four major diseases: **(A)** autoimmune diseases with targets in circulation (circulation-soluble, [CS]) (P = 0.1, Pearson's correlation), **(B)** hematologic malignancies (circulation-membranous, [CM]) (P = 0.9, Pearson's correlation), **(C)** autoimmune diseases with tissue targets (tissue-soluble, [TS]) (P = 0.2, Pearson's correlation), and **(D)** solid tumors (tissue-membranous, [TM]) (P = 0.2, Pearson's correlation).

**Supplementary Figure 2**

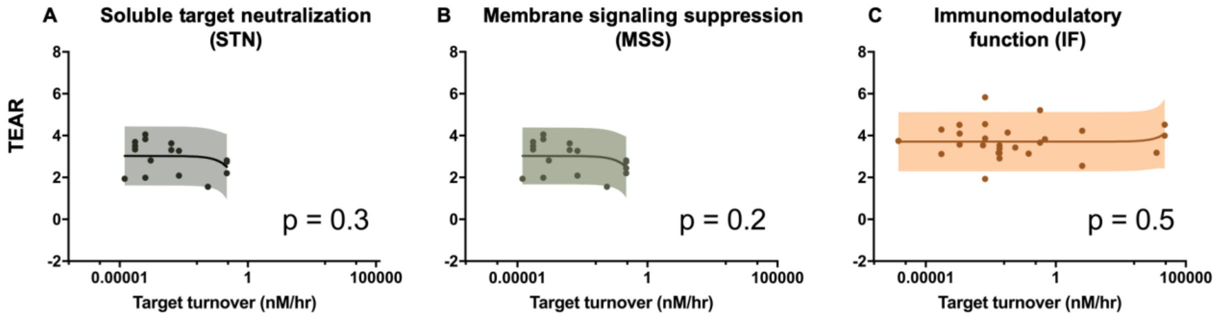


**Supplementary Figure 2. Target turnovers are not relevant to TEARs in three mechanism of action groups.** Dots represent the mean values. Horizontal bars represent SD in the target turnover rates. Vertical bars represent SD in the TEARs. The shadows represent the 90% prediction intervals. No significant correlation between target turnovers and TEARs was observed in three mechanism of action groups: **(A)** soluble target neutralizing (STN) (P = 0.3, Pearson's correlation), **(B)** membranous signaling suppression (MSS) (P = 0.2, Pearson's correlation), and **(C)** immunomodulatory function (IF) (P = 0.5, Pearson's correlation).

**Supplementary Figure 3**

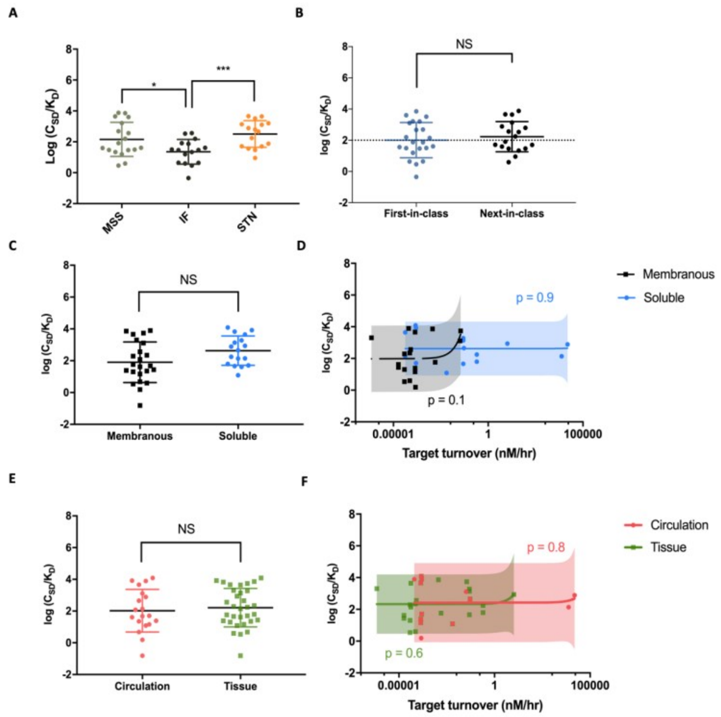


**Supplementary Figure 3. Effects of mechanisms of actions, antibody development, target locations, forms, and turnovers on first-in-human doses (FIHDs). (A)** The log ($C_{FIHD}/K_D$) values are significantly different between antibodies with varying mechanisms of action. The log ($C_{FIHD}/K_D$) values in the MSS and STN groups are significantly higher than the IF group (P = 0.02, P = 0.0005, unpaired Student's *t*-test). **(B)** The log ($C_{FIHD}/K_D$) values are not significantly different between the first-class and the next-class antibodies (P = 0.5, unpaired Student's *t*-test). **(C)** The log ($C_{FIHD}/K_D$) values are not significantly different between the soluble and membranous groups (P = 0.05, unpaired Student's *t*-test). **(D)** Target turnover is not a significant factor to FIHDs in both the membranous and soluble groups. The log ($C_{FIHD}/K_D$) values have no significant correlation with target turnovers, regardless of the solubility of the target (P = 0.1, membranous targets; P = 0.9, soluble targets. Pearson's correlation). **(E)** Target anatomical location does not have a significant impact on FIHs. There is no significant difference between the log ($C_{FIHD}/K_D$) values in the circulation and tissue groups (P = 0.6, unpaired Student's *t*-test). **(F)** Target turnovers are not relevant to antibody FIHDs, either in the circulation group or the tissue group (P = 0.8, P = 0.6, respectively, Pearson's correlation). In **(A), (B), (C)**, and **(E)**, Each dot represents the mean log ($C_{FIHD}/K_D$) values of an antibody. The data is represented in mean ± SD. In **(D)** and **(F)**, dots represent the mean value of an antibody. Horizontal bars represent SD in the target turnover rates. Vertical bars represent SD in the log ($C_{FIHD}/K_D$) values. The shadows represent the 90% prediction intervals.

### Supplementary Table 1

*Supplementary Table 1. Summary of antibody dosing regimens, PK information, targets, mode of actions, and applications*

| Antibody | Maintenance Dose | | | F | $K_D$ | CL | | TEAR | Target | MoA | Applications | Ref |
|---|---|---|---|---|---|---|---|---|---|---|---|---|
| Abatacept | 750 | mg Q4W | IV | | 6.66; 0.361 | 0.28 | mL/h/kg[†a] | 2.60 (0.90) | CD80/CD86 | MSS | Adult rheumatoid arthritis, juvenile idiopathic arthritis | [1, 2] |
| Adalimumab | 40 | mg Q2W | SC | 0.64 | 0.0086; 0.0304; 0.1 | 12 | mL/h/70kg[†a] | 3.16 (0.53) | TNF-α | STN | Rheumatoid arthritis, juvenile idiopathic arthritis, psoriatic arthritis, ankylosing spondylitis, Crohn's disease, plague psoriasis | [3-6] |
| Alirocumab | 150 | mg Q2W | SC | 0.85 | 0.58 | 0.0124 | L/h/70kg[‡a] | 2.56 | PCSK9 | STN | Hypercholesterolemia | [7-9] |
| Atezolizumab | 1200 | mg Q3W | IV | | 0.4 | 0.2 | L/d/70kg[†a] | 3.69 | PD-L1 | MSS/IF | Locally advanced or metastatic urothelial carcinoma, metastatic non-small cell lung cancer | [10, 11] |
| Avelumab | 10 | mg/kg Q2W | IV | | 0.7 | 0.59 | L/d/70kg[†a] | 2.92 | PD-L1 | MSS/IF | Metastatic Merkel cell carcinoma | [12-14] |
| Belatacept | 5 | mg/kg Q4W | IV | | 0.423; 0.035 | 0.51 | mL/h/kg[†a] | 3.12 (0.77) | CD80/CD86 | MSS | Immunosuppression | [2, 15] |
| Belimumab | 10 | mg/kg Q4W | IV | | 0.25-0.35 | 215 | mL/d/70kg[†a] | 3.42 (0.10) | BAFF | STN | Systemic lupus erythematosus | [16, 17] |
| Benralizumab | 30 | mg Q8W | SC | 0.58 | 0.011 | 0.29 | L/d/70kg[†a] | 2.81 | IL-5R α | MSS | Asthma | [18] |
| Bevacizumab | 10 | mg/kg Q2W | IV | | 1.1 | 0.207 | L/d/70kg[†a] | 3.17 | VEGF | STN | Cervical cancer, glioblastoma, metastatic colorectal cancer, metastatic renal cell carcinoma, non-squamous non-small cell lung cancer | [19, 20] |
| Brodalumab | 210 | mg Q2W | SC | 0.55 | 0.239 | 3 | L/d/70kg[‡b] | 2.15 | IL-17RA | MSS | Plaque psoriasis | [21, 22] |
| Burosumab | 1 | mg/kg Q4W | SC | 0.9 | 0.01 | 0.29 | L/d/70kg[†b] | 3.77 | FGF23 | STN | X-linked hypophosphatemia | [23, 24] |

[†]Linear systemic clearance at the maintenance dose; [‡]Non-linear systemic clearance at the maintenance dose; [a] CL; [b] CL/F; F: bioavailability; MoA = Mechanism of Action; IV = Intravenous injection; SC = Subcutaneous injection; IM= intramuscular injection; STN = Soluble Target Neutralization; MSS= Membrane Signaling Suppression; IF= Immunomodulatory Functions; TEAR = Therapeutic Exposure Affinity Ratio.

*Supplementary Table 1. Summary of antibody dosing regimens, PK information, targets, mode of actions, and applications*

| Antibody | Maintenance Dose | | | F | $K_D$ | CLp | | TEAR | Target | MoA | Applications | Ref |
|---|---|---|---|---|---|---|---|---|---|---|---|---|
| Canakinumab | 150 | mg Q8W | SC | 0.66 | 0.0305 | 0.174 | L/d/70kg[†a] | 3.36 | IL-1β | STN | Periodic fever syndromes, active systemic juvenile idiopathic arthritis | 25-27 |
| Cemiplimab | 350 | mg Q3W | IV | | 5.61 | 0.21 | L/d/70kg[†a] | 1.99 | PD-1 | MSS/IF | Metastatic cutaneous squamous cell carcinoma | 28, 29 |
| Cetuximab | 250 | $mg/m^2$ QW | IV | | 0.39 | 0.497 | L/day/70kg[‡a] | 3.31 | EGFR | MSS/IF | Head and neck cancer, colorectal cancer | 30-32 |
| Daratumumab | 16 | mg/kg Q4W | IV | | 4.36 | 171.4 | mL/d/70kg[‡a] | 2.56 | CD38 | IF | Multiple myeloma | 33, 34 |
| Denosumab | 60 | mg Q6M | SC | 0.61 | 0.003 | 3.25 | mL/h/66kg[‡a] | 3.75 | RANKL | MSS/STN | Osteoporosis, increase bone mass | 35-37 |
| Dinutuximab | 17.5 | $mg/m^2 \times 4$ Q28D | IV | | 11.2 | 0.21 | L/d/70kg[†a] | 1.09 | GD2 | IF | Neuroblastoma | 38 |
| Dupilumab | 300 | mg Q2W | SC | 0.6 | 0.03; 0.012 | 0.126 | L/d/70kg[‡a] | 4.56 (0.28) | IL-4R α | MSS | Atopic dermatitis | 39-41 |
| Durvalumab | 10 | mg/kg Q2W | IV | | 0.667 | 0.232 | L/d/70kg[†a] | 3.34 | PD-L1 | MSS/IF | Urothelial carcinoma, non-small cell lung cancer | 13, 42, 43 |
| Eculizumab | 1200 | mg Q2W | IV | | 0.05 | 14.6 | ml/hr/70kg[†a] | 4.52 | C5 | STN | Atypical hemolytic uremic syndrome, paroxysmal nocturnal hemoglobinuria | 44, 45 |
| Elotuzumab | 10 | mg/kg Q2W | IV | | 43.7 | 0.0895 | L/day/70kg[‡a] | 1.94 | SLAMF7 | IF | Multiple myeloma | 46-48 |
| Emapalumab | 1 | mg/kg Q3D | IV | | 0.0014 | 0.007 | L/h/70kg[†a] | 5.83 | IFNγ | STN | Hemophagocytic lymphohistiocytosis | 49, 50 |
| Erenumab | 140 | mg QMT | SC | 0.82 | 0.02 | 0.214 | L/d/70kg[‡a] | 3.78 | CGRP-R | MSS | Migraine | 51, 52 |

[†]Linear systemic clearance at the maintenance dose; [‡]Non-linear systemic clearance at the maintenance dose; [a] CL; [b] CL/F; F: bioavailability; MoA = Mechanism of Action; IV = Intravenous injection; SC = Subcutaneous injection; IM= intramuscular injection; STN = Soluble Target Neutralization; MSS= Membrane Signaling Suppression; IF= Immunomodulatory Functions; TEAR = Therapeutic Exposure Affinity Ratio.

*Supplementary Table 1. Summary of antibody dosing regimens, PK information, targets, mode of actions, and applications*

| Antibody | Maintenance Dose | | | F | $K_D$ | CLp | | TEAR | Target | MoA | Applications | Ref |
|---|---|---|---|---|---|---|---|---|---|---|---|---|
| Etanercept | 50 | mg QW | SC | 0.58 | 0.0004, 0.4 | 132 | ml/hr/70kg$^{\dagger b}$ | 3.07 (2.12) | TNF | STN | Rheumatoid arthritis, polyarticular juvenile idiopathic arthritis, psoriatic arthritis, ankylosing spondylitis, plaque psoriasis | 4, 53-55 |
| Evolocumab | 420 | mg QM | SC | 0.72 | 0.016 | 0.256 | L/d/70kg$^{\ddagger a}$ | 4.23 | PCSK9 | STN | Hypercholesterolemia | 56-58 |
| Fremanezumab | 225 | mg QM | SC | 0.66 | 0.0022 | 0.141 | L/d/70kg$^{\dagger b}$ | 5.21 | CGRP ligand | STN | Migraine | 59-61 |
| Galcanezumab | 120 | mg QMT | SC | 0.82 | 0.031 | 0.008 | L/hr/70kg$^{\dagger b}$ | 3.66 | CGRP ligand | STN | Migraine | 62-64 |
| Golimumab | 50 | mg QM | SC | 0.53 | 0.018 | 0.4 | L/d/70kg$^{\dagger a}$ | 2.91 | TNF-α | MSS/STN | Rheumatoid arthritis, active psoriatic arthritis, ankylosing spondylitis | 65-67 |
| Guselkumab | 100 | mg Q8W | SC | 0.49 | 0.0033 | 0.516 | L/d/70kg$^{\dagger b}$ | 3.86 | IL-23 | STN | Plaque psoriasis | 68, 69 |
| Ibalizumab | 800 | mg Q2W | IV | | 0.082 | 0.0121 | L/hr/70kg$^{\ddagger a}$ | 4.20 | CD4 | MSS | HIV infection | 70-72 |
| Infliximab | 5 | mg/kg Q6W | IV | | 0.0042; 0.117; 0.45; 0.027 | 0.23; 0.37; 0.41; 0.29; 0.38 | L/d/70kg$^{\dagger a}$ | 3.53 (0.79) | TNF-α | MSS/STN | Crohn's disease, ulcerative colitis, rheumatoid arthritis, ankylosing spondylitis, psoriatic arthritis, plaque psoriasis | 4, 5, 73, 74 |
| Ipilimumab | 3 | mg/kg Q3W | IV | | 5.25 | 14.9 | ml/hr/70kg$^{\dagger a}$ | 1.56 | CTLA-4 | MSS/IF | Metastatic melanoma, cutaneous melanoma | 75-77 |
| Ixekizumab | 80 | mg Q4W | SC | 0.6 – 0.81 | 0.0018 | 0.39 | L/d/70kg$^{\dagger a}$ | 4.29 | IL-17A | STN | Plaque psoriasis, psoriatic arthritis | 78, 79 |
| Lanadelumab | 300 | mg Q2W | SC | 0.66 | 0.12 | 0.809 | L/d/70kg$^{\dagger b}$ | 3.18 | plasma kallikrein | STN | Hereditary angioedema prevention | 80, 81 |
| Mepolizumab | 300 | mg Q4W | SC | 0.8 | 0.0042, 0.1 | 0.28 | L/d/70kg$^{\dagger b}$ | 4.10 (0.97) | IL-5 | STN | Asthma, eosinophilic granulomatosis | 82, 83 |

$^{\dagger}$Linear systemic clearance at the maintenance dose; $^{\ddagger}$Non-linear systemic clearance at the maintenance dose; $^{a}$ CL; $^{b}$ CL/F; F: bioavailability; MoA = Mechanism of Action; IV = Intravenous injection; SC = Subcutaneous injection; IM= intramuscular injection; STN = Soluble Target Neutralization; MSS= Membrane Signaling Suppression; IF= Immunomodulatory Functions; TEAR = Therapeutic Exposure Affinity Ratio.

*Supplementary Table 1. Summary of antibody dosing regimens, PK information, targets, mode of actions, and applications*

| Antibody | Maintenance Dose | | | F | $K_D$ | CLp | | TEAR | Target | MoA | Applications | Ref |
|---|---|---|---|---|---|---|---|---|---|---|---|---|
| Natalizumab | 300 | mg Q4W | IV | | 0.3 | 16 | ml/hr/70kg[‡a] | 2.79 | α4β1/α4β7 | MSS | Multiple sclerosis, Crohn's disease | 84, 85 |
| Necitumumab | 800 | mg D1 + D8 Q3W | IV | | 0.32 | 14.1 | ml/hr/70kg[†a] | 3.63 | EGFR | MSS/IF | Metastatic squamous non-small cell lung cancer | 86, 87 |
| Nivolumab | 240 | mg Q2W | IV | | 2.6 | 8.2 | ml/hr/70kg[†a] | 2.36 | PD1 | MSS/IF | Metastatic melanoma, metastatic non-small cell lung cancer, advanced renal cell carcinoma, Hodgkin lymphoma, metastatic squamous cell carcinoma, metastatic urothelial carcinoma, hepatocellular carcinoma | 88, 89 |
| Obinutuzumab | 1000 | mg Q28D | IV | | 4 | 0.11, 0.08 | L/d/70kg[†a] | 2.81 (0.10) | CD20 | IF | Chronic lymphoid leukemia, follicular lymphoma | 90, 91 |
| Ocrelizumab | 600 | mg Q6M | IV | | 0.84 | 0.17 | L/d/70kg[†a] | 2.21 | CD20 | IF | Multiple sclerosis | 92, 93 |
| Ofatumumab | 2000 | mg Q4W | IV | | 4.76 | 7.5 | ml/hr/70kg[‡a] | 2.75 | CD20 | IF | Chronic lymphoid leukemia | 94-96 |
| Olaratumab | 15 | mg/kg D1 + D8 Q21D | IV | | 0.04; 0.33 | 0.56 | L/d/70kg[†] | 4.00 (0.65) | PDGFR-α | MSS | Soft tissue sarcoma | 97-100 |
| Omalizumab | 375 | mg Q4W | SC | 0.62 | 0.02 – 7.7 | 2.4 | ml/d/kg[†b] | 3.13 (1.83) | IgE | STN | Chronic idiopathic urticaria, asthma | 101, 102 |
| Palivizumab | 15 | mg/kg QM | IM | 0.7 | 0.96 | 198 | ml/d/70kg[†a] | 2.97 | RSV | STN | Prevention of respiratory tract disease caused by respiratory syncytial virus | 103-105 |
| Panitumumab | 6 | mg/kg Q14D | IV | | 0.05 | 4.9 | mL/d/kg[‡a] | 4.08 | EGFR | MSS | Metastatic colorectal cancer | 30, 106, 107 |

[†]Linear systemic clearance at the maintenance dose; [‡]Non-linear systemic clearance at the maintenance dose; [a] CL; [b] CL/F; F: bioavailability; MoA = Mechanism of Action; IV = Intravenous injection; SC = Subcutaneous injection; IM= intramuscular injection; STN = Soluble Target Neutralization; MSS= Membrane Signaling Suppression; IF= Immunomodulatory Functions; TEAR = Therapeutic Exposure Affinity Ratio.

*Supplementary Table 1. Summary of antibody dosing regimens, PK information, targets, mode of actions, and applications*

| Antibodiy | Maintenance Dose | | | F | $K_D$ | CLp | | TEAR | Target | MoA | Applications | Ref |
|---|---|---|---|---|---|---|---|---|---|---|---|---|
| Pembrolizumab | 200 | mg Q3W | IV | | 0.029 | 195 | mL/d/70kg[†a] | 4.05 | PD-1 | MSS/IF | Melanoma, non-small cell lung cancer, head and neck squamous cell cancer, classical Hodgkin lymphoma, primary mediastinal large B-cell lymphoma, urothelial carcinoma, microsatellite instability-high cancer, etc | 108-110 |
| Pertuzumab | 420 | mg Q3W | IV | | 0.3 | 0.24 | L/d/70kg[†a] | 3.27 | HER2 | MSS/IF | HER2-positive metastatic breast cancer | 111, 112 |
| Ramucirumab | 8 | mg/kg Q2W | IV | | 0.05 | 0.015 | L/hr/70kg[‡a] | 4.18 | VEGFR2 | MSS | Advanced gastric or gastro-esophageal junction adenocarcinoma, metastatic nonsmall cell lung cancer, metastatic colorectal cancer | 113, 114 |
| Ravulizumab | 3300 | mg Q8W | IV | | 0.5 | 0.08 | L/d/70kg[†a] | 4.00 | C5 | STN | Paroxysmal nocturnal hemoglobinuria | 115, 116 |
| Reslizumab | 3 | mg/kg Q4W | IV | | 0.081 | 7 | ml/hr/70kg[†a] | 3.57 | IL-5 | STN | Asthma | 117, 118 |
| Risankizumab | 150 | mg Q12W | SC | 0.89 | 0.001 | 0.31 | L/d/70kg | 4.55 | IL-23 | STN | Plaque psoriasis | 119-121 |
| Rituximab | 375 | $mg/m^2$ QW | IV | | 5.2 – 11 | 0.335; 0.312; 0.252 | L/d/70kg[‡a] | 2.45 (0.19) | CD20 | IF | Non-Hodgkin's lymphoma, chronic lymphocytic leukemia, rheumatoid arthritis, granulomatosis with polyangiitis, microscopic polyangiitis | 122-124 |
| Romosozumab | 210 | mg QM | SC | 0.81 | 0.00063; 0.0063 | 0.38 | mL/hr/kg[‡b] | 4.57 (0.71) | Sclerostin | STN | Osteoporosis | 125-127 |

[†]Linear systemic clearance at the maintenance dose; [‡]Non-linear systemic clearance at the maintenance dose; [a] CL; [b] CL/F; F: bioavailability; MoA = Mechanism of Action; IV = Intravenous injection; SC = Subcutaneous injection; IM= intramuscular injection; STN = Soluble Target Neutralization; MSS= Membrane Signaling Suppression; IF= Immunomodulatory Functions; TEAR = Therapeutic Exposure Affinity Ratio.

*Supplementary Table 1. Summary of antibody dosing regimens, PK information, targets, mode of actions, and applications*

| Antibodiy | Maintenance Dose | | | F | $K_D$ | CLp | | TEAR | Target | MoA | Applications | Ref |
|---|---|---|---|---|---|---|---|---|---|---|---|---|
| Sarilumab | 200 | mg Q2W | SC | 0.8 | 0.054 | 0.26 | L/d/70kg[‡b] | 3.83 | IL-6R | MSS/ STN | Rheumatoid arthritis | 128-130 |
| Secukinumab | 300 | mg Q4W | SC | 0.55 – 0.77 | 0.2 | 0.14 – 0.22 | L/d/70kg[†a] | 3.12 (0.14) | IL-17A | STN | Plaque psoriasis, psoriatic arthritis, ankylosing spondylitis | 131, 132 |
| Siltuximab | 11 | mg/kg Q3W | IV | | 0.034 | 0.23 | L/d/70kg[†a] | 4.51 | IL-6 | MSS/ STN | Multicentric Castleman's disease | |
| Tildrakizumab | 100 | mg Q12W | SC | 0.73 – 0.8 | 0.297 | 0.32 | L/d/70kg[†a] | 1.82 | IL-23 | STN | Plaque psoriasis | 133, 134 |
| Tocilizumab | 8 | mg/kg Q4W | IV | | 2.54 | 0.3 | L/d/70kg[‡a] | 2.25 | IL-6R | MSS | Rheumatoid arthritis, giant cell arteritis, polyarticular juvenile idiopathic arthritis, systemic juvenile idiopathic arthritis, cytokine release syndrome | 135-137 |
| Trastuzumab | 2 | mg/kg QW | IV | | 5 | 0.225 | L/d/70kg[†a] | 2.09 | HER2 | MSS/I F | HER2-overexpressing breast cancer, HER2-overexpressing metastatic gastric cancer | 138-140 |
| Vedolizumab | 300 | mg Q8W | IV | | 0.47 | 0.157 | L/d/70kg[‡a] | 2.69 | α4β7 integrin | MSS | Adult ulcerative colitis, adult Crohn's disease | 141, 142 |

[†]Linear systemic clearance at the maintenance dose; [‡]Non-linear systemic clearance at the maintenance dose; [a] CL; [b] CL/F; F: bioavailability; MoA = Mechanism of Action; IV = Intravenous injection; SC = Subcutaneous injection; IM= intramuscular injection; STN = Soluble Target Neutralization; MSS= Membrane Signaling Suppression; IF= Immunomodulatory Functions; TEAR = Therapeutic Exposure Affinity Ratio.

### Supplementary Table 2

*Supplementary Table 2. Target information of the included therapeutic antibodies.*

| Target | Plasma baseline | | Half life | | Target turnover | Ref |
|---|---|---|---|---|---|---|
| BAFF | 1.5 | ng/mL | 70 | min | 27.84 | 143, 144 |
| C5 | 0.37 | uM | 1 | min | 15384600 | 145, 146 |
| CD20 | 200000 | molecules/cell | 426 | hr | 0.15 | 147 |
| CD4 | 46,000 | molecules/cell | 35 | min | 1270.88 | 148-150 |
| CD80 | 0.13, 0.29, 0.28 | ng/ml | 14000 | s | 0.73 (0.23) | 151, 152 |
| CD86 | 1.46, 1.85, 1.71, 1.76 | ng/ml | 5† | hr | 3.36 (0.33) | 152 |
| CGRP ligand | 42 | pmol/L | 7 | min | 249.48 | 153, 154 |
| CGRP receptor | 2796 | molecules/cell | 0.98 | day | 0.039 | 155, 156 |
| CTLA-4 | 1.61, 3.19, 4.05, 3.90 | ng/ml | 151 | min | 27.6 (9.16) | 152, 157 |
| EGFR | 50000 | molecules/cell | 8-24 | hr | 1.03 | 158, 159 |
| FGF23 | 26.1 | pg/ml | 20-40 | min | 1.61 | 160, 161 |
| GD2 | 10000000 | molecules/cell | 114, 462 | min | 1082.11 (926.25) | 162, 163 |
| HER2 | 100000 | molecules/cell | 8 | hr | 2.06 | 164, 165 |
| IFNγ | 139.6, 179, 151 | pg/ml | 94 | min | 1.54 (0.20) | 166, 167 |
| IgE | 368; 1123;1682; 2529 | ng/ml | 2.4 | day | 90.27 (57.73) | 168, 169 |
| IL-17 α | 200; 20; 8.29 | pg/ml | 24-48 | hr | 0.042 (0.059) | 170-172 |
| IL-1$\beta$ | 1 | ng/ml | 1.59 | hr | 14.17 | 173, 174 |
| IL-23 | 477; 799 | pg/ml | 4 | hr | 1.93 (0.69) | 175, 176 |
| IL-4R α | 25.6 (s) | pg/ml | 80 – 90 | min | 0.089 | 175, 177, 178 |
| IL-5 | 26; 15 | pg/ml | 6 | hr | 0.16 (0.06) | 179, 180 |
| IL-5R α | 42.3 (s) | pg/ml | 3 | hr | 0.16 | 181, 182 |
| IL-6 | 10 | pg/ml | 0.75; 1.7; 6 | hr | 0.20 (0.17) | 170, 183, 184 |
| IL-6R | 58000 | pg/mL | 2-3 | hr | 380.53 | 170, 185, 186 |
| PCSK9 | 89.5 | ng/ml | 5 | min | 10337.25 | 187, 188 |
| PD-L1 | 42.21; 37.81 | pg/mL | 18 | hr | 0.039 (0.0030) | 189, 190 |
| PD1 | 2.9 | ng/mL | 281.7 | hr | 0.14 | 191, 192 |
| PDGFRα | 9914 | molecules/cell | 3 | min | 65.22 | 193, 194 |
| Plasma kallikrein | 70-90 | ug/ml | 5 | min | 7560000 | 195, 196 |
| RANKL | 0.6 | pM | 468 | hr | 0.00089 | 197, 198 |
| SLAMF7 | 0.255 | ng/ml | 11 | day | 0.016 | 46, 199 |
| TNF-alpha | 3.27; 7.79; 1000; 170 | pg/mL | 70 | min | 6.85 (11.05) | 200-203 |
| VEGF | 75; 30 | pg/mL | 0.7; 34 | min | 6.39 (5.25) | 204-207 |
| VEGFR2 | 12794, 13625 | pg/mL | 70 | min | 35.26 (1.57) | 208, 209 |

†Estimated value. The target turnover was presented as mean $\pm$ SD. The unit of target turnover was $10^{-3}$ nM/hr.

**Supplementary Table 3**

*Table 3. Summary of the first-in-human (FIH) doses, MAD (Maximum Administered Doses), and P2D (Phase II doses) of surveyed antibodies*

| | FIH dose | | log ($C_{FIHD}/K_D$) | MAD | | log ($C_{FIHD}/K_D$) | P2D | | log ($C_{P2D}/K_D$) | P2D Selection | Ref |
|---|---|---|---|---|---|---|---|---|---|---|---|
| Atezolizumab[1] | 0.01 | mg/kg Q3W IV | 0.46 | 20 | mg/kg Q3W IV | 3.76 | 1200[a] | mg Q3W IV | 3.69 | MABEL | 11, 210 |
| Avelumab[2] | 1 | mg/kg Q2W IV | 1.91 | 20 | mg/kg Q2W IV | 3.21 | 10[a] | mg/kg Q2W IV | 2.92 | MABEL | 211, 212 |
| Belimumab[1] | 1 | mg/kg single IV | 2.72 | 20 | mg/kg Q3W IV | 3.85 | 300 | mg QW SC | 3.66 | NA | 213, 214 |
| Benralizumab[2] | 0.03 | mg/kg single IV | 2.50 | 3 | mg/kg single IV | 4.50 | 100 | mg Q8W SC | 3.57 | Efficacy | 215-217 |
| Bevacizumab[1] | 0.1 | mg/kg Q28D IV | 0.85 | 10 | mg/kg Q28D IV | 2.85 | 20 | mg/kg Q2W IV | 3.17 | MABEL | 218, 219 |
| Brodalumab[1] | 7 | mg single SC | 0.69 | 700 | mg single IV | 2.65 | 210 | mg Q2W SC | 2.17 | Efficacy | 220 |
| Burosumab[1] | 0.003 | mg/kg, single IV | 1.59 | 1 | mg/kg single SC | 3.77 | 2 | mg/kg Q2W SC | 4.07 | Efficacy | 221, 222 |
| Cemiplimab[2] | 1 | mg/kg Q2W IV | 1.46 | 10 | mg/kg Q2W IV | 2.46 | 350 | mg Q3W IV | 1.99 | NA | 223 |
| Cetuximab[1] | 5 | $mg/m^2$ single IV | 1.78 | 400 | $mg/m^2$ QW IV | 3.99 | 250 | $mg/m^2$ QW IV | 3.78 | MABLE | 31, 224 |
| Daratumumab[1] | 0.005 | mg/kg QW IV | -0.35 | 24 | mg/kg QW IV | 3.33 | 16 | mg/kg Q4W IV | 2.56 | MABEL | 225, 226 |
| Dupilumab[1] | 1 | mg/kg single IV | 3.95 | 12 | mg/kg single IV | 5.03 | 300 | mg QW SC | 4.70 | Efficacy | 227, 228 |
| Durvalumab[2] | 0.1 | mg/kg Q2W IV | 1.34 | 10 | mg/kg Q2W IV | 3.34 | 10[a] | mg/kg Q2W IV | 3.34 | MABEL | 229 |
| Elotuzumab[1] | 0.5 | mg/kg Q2W IV | 0.63 | 20 | mg/kg Q2W IV | 2.23 | 20 | mg/kg QW IV | 2.53 | MABEL | 230, 231 |
| Erenumab[1] | 21 | mg single SC | 3.28 | 210 | mg single SC | 4.28 | 140 | mg Q4W SC | 3.81 | MABEL | 232, 233 |
| Evolocumab[2] | 7 | mg single SC | 2.79 | 420 | mg single IV | 4.41 | 420 | mg Q4W SC | 4.26 | Efficacy | 234 |
| Fremanezumab[2] | 0.2 | mg single IV | 2.49 | 2000 | mg single IV | 6.49 | 900 | mg Q28D SC | 5.85 | Efficacy | 235, 236 |
| Galcanezumab[2] | 1 | mg single SC | 1.91 | 600 | mg single SC | 4.69 | 150[a] | mg Q2W SC | 4.09 | Efficacy | 62 |
| Guselkumab[1] | 0.03 | mg/kg single IV | 3.53 | 10 | mg/kg single IV | 6.05 | 200 | mg Q12W SC | 4.73 | Efficacy | 69, 237 |
| Ibalizumab[1] | 0.3 | mg/kg single IV | 3.23 | 25 | mg/kg single IV | 5.15 | 25 | mg/kg Q2W IV | 5.15 | Efficacy | 238, 239 |
| Ixekizumab[2] | 0.06 | mg/kg single IV | 3.45 | 2 | mg/kg single IV | 4.98 | 150 | mg QM SC | 4.55 | Efficacy | 240, 241 |
| Lanadelumab[1] | 0.1 | mg/kg single SC | 1.63 | 3 | mg/kg single SC | 3.10 | 400 | mg Q14D SC | 3.38 | Efficacy | 242, 243 |

[1] First-in-class antibodies; [2] second-in-class antibodies; [a] Recommended Phase II dose; IV = Intravenous injection; SC = Subcutaneous injection; FIH = First-in-Human; MAD = Maximum Administered Dose in FIH trials; P2D = Phase II Doses; MABEL = Minimum Anticipated Biological Effect Level; NA = Not Available.

*Table 3. Summary of the first-in-human (FIH) doses, MAD (Maximum Administered Doses), and P2D (Phase II doses) of surveyed antibodies*

| | FIH dose | | log ($C_{FIHD}/K_D$) | MAD | | log ($C_{FIHD}/K_D$) | P2D | | log ($C_{P2D}/K_D$) | P2D Selection | Ref |
|---|---|---|---|---|---|---|---|---|---|---|---|
| Mepolizumab[1] | 0.5 | mg/kg single IV | 3.88 | 10 | mg/kg single IV | 5.18 | 750 | mg Q4W IV | 5.18 | Efficacy | 244, 245 |
| Nivolumab[1] | 0.3 | mg/kg single IV | 2.75 | 10 | mg/kg single IV | 4.28 | 3 | mg/kg Q2W IV | 2.30 | Efficacy/MABEL | 246, 247 |
| Obinotuzumab[1] | 100 | mg Q21D IV | 1.93 | 2000 | mg Q21D IV | 3.23 | 800 [a] | mg Q3W IV | 2.84 | MABEL | 248, 249 |
| Ocrelizumab[1] | 10 | mg Q15D IV | 1.51 | 1000 | mg Q15D IV | 3.51 | 2000 | mg Q24W IV | 2.74 | Efficacy | 250, 251 |
| Ofatumumab[2] | 300 | mg QW IV | 2.53 | 1000 | mg QW IV | 3.05 | 1000 [a] | mg QW IV | 3.05 | Efficacy | 252, 253 |
| Olaratumab[1] | 4 | mg/kg QW IV | 3.81 | 16 | mg/kg QW IV | 4.42 | 16 [a] | mg/kg QW IV | 4.42 | Efficacy | 254 |
| Panitumumab[2] | 0.01 | mg/kg QW IV | 1.60 | 5 | mg/kg QW IV | 4.30 | 6 | mg/kg Q2W IV | 4.08 | MABEL | 255, 256 |
| Pembrolizumab[2] | 1 | mg/kg Q2W IV | 3.77 | 10 | mg/kg Q2W IV | 4.77 | 2 [a] | mg/kg Q3W IV | 3.90 | MABEL | 257 |
| Pertuzumab[1] | 0.5 | mg/kg Q3W IV | 2.19 | 15 | mg/kg Q3W | 3.67 | 420 [a] | mg Q3W | 3.27 | Efficacy | 258, 259 |
| Ramucirumab[2] | 2 | mg/kg QW IV | 3.88 | 16 | mg/kg QW IV | 4.78 | 8 [a] | mg/kg QW IV | 4.48 | MABEL | 260, 261 |
| Ravulizumab[2] | 200 | mg single IV | 3.38 | 400 | mg single IV | 3.68 | 5400 | mg Q12W IV | 4.04 | MABEL | 262, 263 |
| Reslizumab[2] | 0.03 | mg/kg single IV | 1.87 | 1 | mg/kg single IV | 3.40 | 3 | mg/kg Q4W IV | 3.57 | Efficacy | 264, 265 |
| Risankizumab[2] | 0.01 | mg/kg single IV | 3.04 | 5 | mg/kg single IV | 5.74 | 180 | mg Q12W SC | 4.63 | Efficacy | 120, 266 |
| Rituximab[1] | 10 | $mg/m^2$ single IV | 0.60 | 500 | $mg/m^2$ single IV | 2.30 | 375 | $mg/m^2$ QW IV | 2.48 | Efficacy | 267, 268 |
| Romosozumab[1] | 0.1 | mg/kg single SC | 3.42 | 10 | mg/kg single SC | 5.16 | 210 | mg QM SC | 4.81 | Efficacy | 269 |
| Siltuximab [2] | 1 | mg/kg Q2W IV | 3.64 | 12 | mg/kg Q2W IV | 4.72 | 15 | mg/kg Q4W IV | 4.52 | Efficacy/MABEL | 270 |
| Tildrakizumab[2] | 0.1 | mg/kg Q56D IV | 0.95 | 10 | mg/kg Q28D | 3.25 | 200 | mg Q12W SC | 2.23 | Efficacy | 271, 272 |
| Trastuzumab[2] | 10 | mg single IV | 0.64 | 500 | mg single IV | 2.34 | 2 | mg/kg QW IV | 2.09 | NA | 273 |

[1] First-in-class antibodies; [2] second-in-class antibodies; [a] Recommended Phase II dose; IV = Intravenous injection; SC = Subcutaneous injection; FIH = First-in-Human; MAD = Maximum Administered Dose in FIH trials; P2D = Phase II Doses; MABEL = Minimum Anticipated Biological Effect Level; NA = Not Available.
